## Supplementary data items and methods for "A mathematical model that predicts human biological age from physiological traits identifies environmental and genetic factors that influence aging": Comput_Methods.docx

**Platform**

All the calculations had been performed in free statistical software “R” (version 3.5.2) on x86_64-apple-darwin15.6.0 (64-bit) platform.

**Phenotype data-vector normalization**

Before inclusion in the model or correlation analysis, each phenotype was first normalized to the mean and divided by its standard deviation. If the phenotype was encoded as a multiple-choice question (for example, do you take naps - often, sometimes, rarely, never?), each answer option was encoded as binary vector (one or zero), and such vectors were normalized the same way. Euclidean distances and Pearson correlations were calculated for each pair of phenotypes.

**Correlation computation**

For evaluation of the correlation between any two vectors, linear regression was performed using the “lm” R function (stats package version 3.6.2). Adjusted R^2^ and p-value had been extracted from such linear models using the “summary(lm())” method. Where applicable, the significance threshold had been adjusted to account for multiple testing using Bonferroni correction.

Since PLS modeling and many other mathematical approaches do not tolerate missing values and in the UKBB dataset that we selected, over 60,000 participants (~15%) lacked at least one measurement for one of the phenotypes, the following mitigation strategies had been used. To avoid excessive imputation, any individual missing more than 15 data points was excluded from the study. In females, the number of selected participants decreased from 222,111 to 215,949 (~2.7% loss), and in males from 188,609 to 183,715 (~2.6% loss). The rest had any missing data imputed using the R package “BiocManager”, function “impute, version 3.8”. Imputation was performed using the “KNN” algorithm.

PLS models were built in R using the package “pls, version 2.7-2”, function “plsr”. All the phenotypes used in the modeling had been normalized to have a mean value of 0, and a standard deviation of 1, before being included in the model. This procedure ensured that the weight of phenotypes would not depend on the units of measurements. The number of components in the PLS model had been selected for each gender as described in the paper. Cross-validation was performed during the model generation with default parameters. Additionally, to test for overfitting, a PLS model had been generated on randomly selected 90% of individuals and tested on the remaining 10% with similar results, suggesting that no overfitting has occurred. An optimal number of components for the PLS model was identified as 11 for males with a minimum root-mean-square error (RMSE) achieved 5.1 years, and 9 for females with a minimum root-mean-square error of 4.9 years. Cross-Validation was performed using 10 segments, and R function “crossvall”. In this implementation, dataset is split into 10 random segments and 10 consecutive models being trained and tested. Each model is trained on 9 segments and tested on the tenth segment, such that each segment serves as a test segment once. Root-mean-square error of prediction is calculated in each instance for each combination of the components. The output of cross-validation is as follows:

For females:

Number of components considered: 9

VALIDATION: RMSEP

Cross-validated using 10 random segments.

(Intercept) 1 comps 2 comps 3 comps 4 comps 5 comps 6 comps

CV	7.917	5.75	5.344	5.052	4.942	4.881	4.847

adjCV	7.917	5.75	5.344	5.052	4.942	4.881	4.846

7 comps 8 comps 9 comps

CV	4.828	4.819	4.815

adjCV	4.828	4.819	4.814

For males:

Number of components considered: 11

VALIDATION: RMSEP

Cross-validated using 10 random segments.

(Intercept) 1 comps 2 comps 3 comps 4 comps 5 comps 6 comps

CV	8.086	5.959	5.542	5.341	5.271	5.202	5.164

adjCV	8.086	5.959	5.542	5.340	5.271	5.202	5.163

7 comps 8 comps 9 comps 10 comps 11 comps

CV	5.142	5.130	5.121	5.114	5.109

adjCV	5.141	5.129	5.120	5.113	5.108

AdjCV is bias-adjusted cross-validation of the error of prediction.

Where appropriate, boundaries of exponential fitting of the mortality data had been defined as 0.99 confidence. Mortality doubling time with age calculated from UKBB participants was very similar to that reported for the UK in other studies.

Genome-wide association studies (GWAS) were performed using linear models separately on males and females. Analysis was performed using Hail-0.2.38 software, running on a Debian Linux cluster with 72 cores. To account for the influence of environmental factors on ∆Age, we corrected for smoking history, education, income level, Townsend deprivation index (a multiparametric measure of socio-economic status), levels of air pollution, and geographic location of the primary house by including these variables as covariates. To account for and correct for the substructure of the population and possible batch effects, we included the first forty (40) principal components derived from genetic data of the participants and a genotyping batch number as covariates. Only people of white British ancestry were used for GWAS analysis. The GWAS results were analyzed using the PheWeb engine (version 1.1.14). The significance of GWAS hits was calculated considering Linkage Disequilibrium (LD) and adjusted for multiple testing. The nearest genes for all identified loci were called using the GRCh37 human reference genome. Furthermore, we excluded questionable SNPs (Supp. table S5) from the analysis due to irregularities in variant frequencies of certain SNPs identified in the UKBB dataset (Kunert-graf et al., 2020), possibly due to their mis-mapping.

During dropout analysis, only top SNP (single SNP in the associated locus with the smallest p-value) was considered. Both effect size (genetic beta), and nominal p-value were reported using the PheWeb engine.

Linkage Disequilibrium (LD) score regression analysis of GWAS results and genetic correlation analysis were performed as described by (Zheng et al., 2017) using Broad University online tool, specifically dedicated to this type of analysis. This analysis indicated that genetic heritability (H^2^) of ∆Age was ~11% (0.108+/-0.009) for females and ~10% (0.096+/-0.008) for males. Additionally, for male ∆Age GWAS, λ_gc_=1.2005, and LD regression intercept= 1.0213+/-0.0083. For female ∆Age GWAS, λ_gc_ =1.2531, and LD regression intercept= 1.0285+/-0.0119.

Specific details for data derivation (or data source) and processing (where applicable) are also described in the text and the corresponding figure captions.
