## Supplementary data items and methods for "A mathematical model that predicts human biological age from physiological traits identifies environmental and genetic factors that influence aging": Supp_info.docx

**Supplemental Item #1**

**Association of computer game playing habits with youthfulness in men and women**

| **2a**  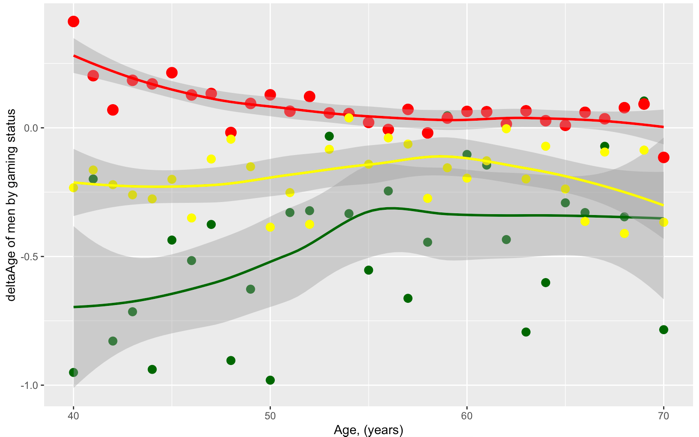 | **2b**  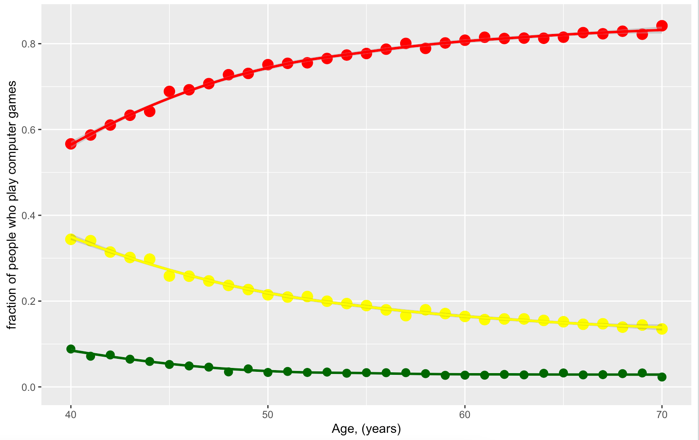 |
| --- | --- |
| **2c**  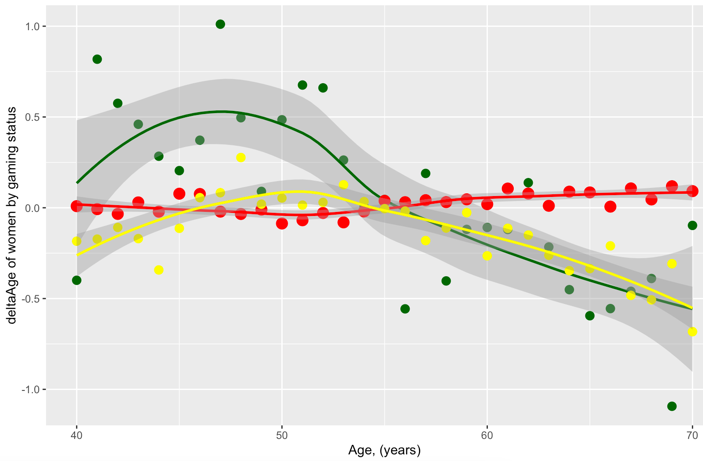 | **2d**  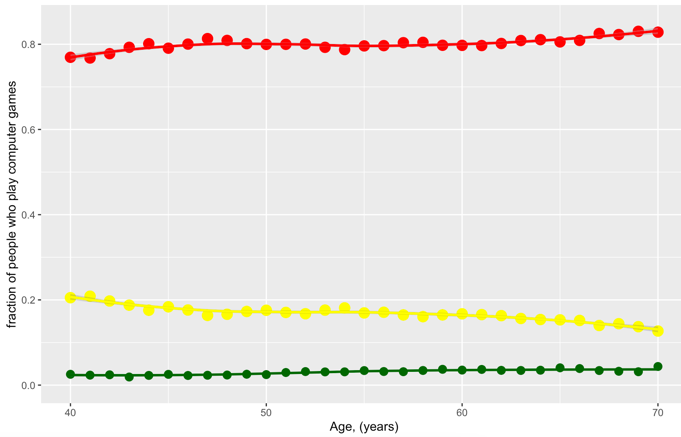 |

Men and older women, who play computer games “often” on average are more youthful. This association is much stronger in men and holds true after age, education and income are considered in the regression. **a)** ∆Age (youthfulness) of males is plotted against age. Three subpopulations are presented – men who play computer games “often” (green dots), men who play computer games “sometimes” (yellow dots), and men who “never” play computer games (red dots). Shaded lines denote 99% confidence intervals; **b)** the fraction of men who play computer games “sometimes”, “never”, or “often” is plotted against age; **c, d)** the same graphs as **a)** and **b)**, except for women. Note that observation is not identical between men and women and is reversed for younger (under the age of 55) women.

**Supplemental Item #2**

**GeneOntology pathways enrichment analysis of GWAS ∆age hits**

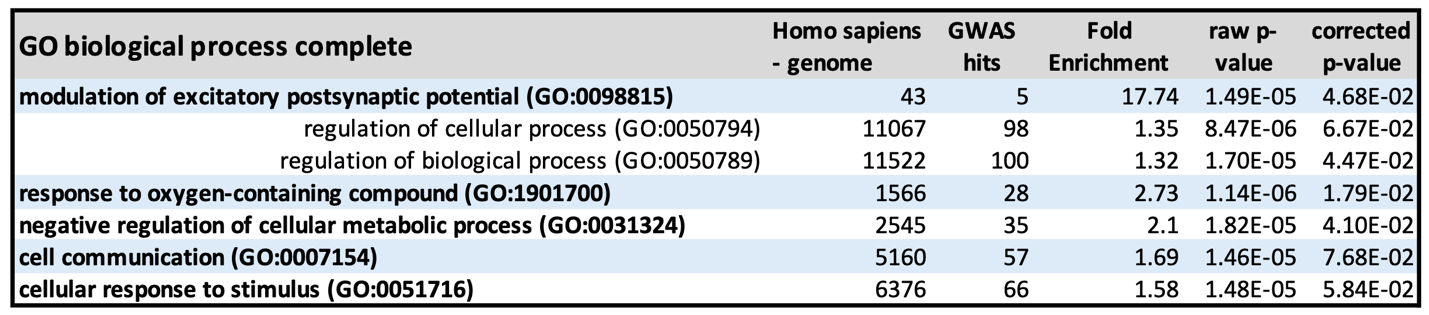

Gene ontology association was performed using an ontology resource previously described ^1^ using an online engine available at <http://geneontology.org/>.
